# Experiments in micro-patterned model membranes support the narrow escape theory

**DOI:** 10.1101/2023.01.03.521408

**Authors:** Elisabeth Meiser, Reza Mohammadi, Nicolas Vogel, David Holcman, Susanne F. Fenz

## Abstract

The narrow escape theory (NET) predicts the escape time distribution of Brownian particles confined to a domain with reflecting borders except for one small window. Applications include molecular activation events in cell biology and biophysics. Specifically, the mean first passage time 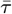 can be analytically calculated from the size of the domain, the escape window, and the diffusion coefficient of the particles. In this study, we systematically tested the NET in a disc by variation of the escape opening. Our model system consisted of micro-patterned lipid bilayers. For the measurement of 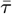, we imaged diffusing fluorescently-labeled lipids using single-molecule fluorescence microscopy. We overcame the lifetime limitation of fluorescent probes by re-scaling the measured time with the fraction of escaped particles. Experiments were complemented by matching stochastic numerical simulations. To conclude, we confirmed the NET prediction *in vitro* and *in silico* for the disc geometry in the limit of small escape openings.

**Significance Statement:** In the biological context of a cell, a multitude of reactions are facilitated by diffusion. It is astonishing how Brownian motion as a cost-efficient but random process is mediating especially fast reactions. The formalism of the narrow escape theory is a tool to determine the average timescale of such processes to be completed (mean first passage time, MFPT) from the reaction space and diffusion coefficient. We present the systematic proof of this formalism experimentally in a bio-mimetic model system and by random walk simulations. Further, we demonstrate a straightforward solution to determine the MFPT from incomplete experimental traces. This will be beneficial for measurements of the MFPT, reliant on fluorescent probes, that have prior been inaccessible.

---

**T**he most economic way for a living cell to reduce energy consumption is to exploit the omnipresent thermal energy. This thermal energy drives Brownian motion at the molecular level (1). However, the random walk of a Brownian particle is not effective in finding a target quickly. Such a target can be a receptor, a channel, or any structure that can trigger a cellular response. In particular, fast activation is critical for stimulus transmission in the nervous system. This leads to the generic question: how can an inefficient stochastic motion mediate a fast response? To overcome this difficulty many cellular processes rely on a large number of copies of identical particles, a situation known as the redundancy principle (2). Another possibility to increase the time performance of a Brownian particle is via adjustment of the cell’s size and shape. As a result, diffusion is often an effective and fast enough mechanism for physiological processes. A theoretical approach to quantify these processes controlled by diffusion in a confining environment with only a small target is the narrow escape theory (NET) (3). In cell biology, there is a multitude of scenarios that can be described by the NET. For example, in the early stages of viral infection, one fundamental step consists in reaching the host cell’s nucleus. To this end, the genetic information of the virus needs to enter through a small nuclear pore (4–6). Another prominent example concerns the activation of postsynaptic receptors by neurotransmitters. Analytical calculations of the mean time of encounter between diffusing transmitters and receptors have revealed the role of receptor distribution in the NET (7).

The present study was motivated by an interesting example of the NET, which is the dynamics of the proteinaceous surface coat of *Trypansoma brucei* (8). *T. brucei* are single-celled eukaryotic parasites and the causative agent of sleeping sickness (9). Trypanosomes exhibit a dense, but dynamic surface coat of glycosylphosphatidylinositol-anchored variant surface glycoproteins (VSG) (10, 11). This coat plays a crucial role in protecting the parasite against the host’s immune system (12). The maintenance of the VSG coat is accomplished by shuffling the VSG through the tiny flagellar pocket, which is the sole site for endo- and exocytosis and makes up only 5 % of the cell surface (13, 14).

The central question of the NET, referred to as the narrow escape problem (NEP), consists in computing the mean first passage time 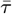 of a Brownian particle until escape through a small exit site on the boundary of a closed domain. The rim of the domain is reflective everywhere except for the exit site, where the particle is absorbed. The mean first passage time characterizes the time scale of rare events. We recall that the concept of the NEP in a cellular context originates from (15) and was extended in (3, 16). The formulas for the NEP were extended to multiple orders in (17, 18). Today, the problem has been well explored by theoreticians (19) in multiple geometries: regular two-dimensional surfaces, cusp geometry or three-dimensional domains connected by narrow cylinders (20).

Here, we will focus on the analytical solution of the NEP for a 2*D*–circular domain, where the mean time 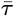 depends predominantly on the diffusion coefficient *D* of the Brownian particle, the area of the disc of radius *r*, and the size of the escape opening *a* (21) :

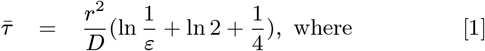

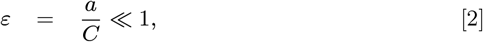

where the parameter *ε* is the relative escape opening defined as the arc length *a* divided by the disk’s circumference *C* = 2*πr*. In the following, the relative escape opening *ε* is referred to as escape opening. Equation 2 is applicable if the Brownian particle starts at least a distance *r* away from the exit site. Experimentally the mean time 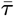 can be estimated by fitting the exponential tail of the empirical probability density histogram of measured escape times (ETs) (20). To our knowledge, two previous studies attempted to measure the mean first passage time *in vitro:* either directly (22) or via Kramer’s rate (23). In the first case, a fluorescent nanoparticle (hydrodynamic diameter of 74 nm) was tracked while diffusing in a small three-dimensional cavity and escaping through a nano-pore. In the second case, the author studied the dynamics of lipid-anchored quantum dots on the membrane of giant unilamellar vesicles escaping onto a lipid nanotube pulled from the vesicle. However, despite the established theory, a systematic experimental verification is still missing.

The critical requirement to measure the mean time 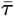 either *in vitro* or *in vivo*, is the reliable traceability of the particles. However, for fluorescent labels, it can easily be hampered by blinking or bleaching (24). Gold beads (25) and nanodiamonds (26, 27) exhibit extremely high photostability and are also non-toxic labels. However, the favorable optical properties are offset by their large size and the disposition to introduce cross-linking of proteins (28). Restraints caused by the labeling can be avoided by using label-free techniques, one example being interferometric scattering microscopy (iSCAT) (29), which, on the downside, is only applicable in *in vitro* experiments.

*In vivo* measurements often rely on fluorescent proteins (30, 31) or smaller organic dyes. The use of fluorescent proteins is especially interesting in live-cell imaging because of the direct assessment of intracellular dynamics, rendering cell permeabilization obsolete. With the increasing size of the fluorescent tag, interference with the folding, localization, and thereby the dynamics of the sample can become a considerable hindrance (32). Organic dyes are in general smaller, ensure minimal interference with the experiment, and are more photo-stable than fluorescent proteins. But even very popular dyes like ATTO647N, which have their emission spectra far apart from the auto-fluorescence of biological samples (33) and provide good photo-stability (34), miss to fully sample escape events in narrow escape scenarios. Especially the extremely long escape times, which lead to the characteristic long tail of the exponential distribution, are out of reach.

In single-molecule fluorescence microscopy, the finite signal-to-noise ratio results in non-continuous tracking. Thus, the achievable trace length is limited further.

The general topic of mortality of Brownian particles was already studied theoretically (20, 35–37). Further, Grebenkov and Rupprecht introduced a comprehensive extension of the narrow escape theory for two and three dimensions, including mortal walkers (38). However, a straightforward solution to overcome lifetime limitations remained elusive for experimentalists. Limitations hampered experimental testing of the NEP. With our work presented here, we aim to fill this gap for the narrow escape problem.

In this study, we propose to test the narrow escape problem experimentally in a disc by varying the size of the escape opening. To this end, we developed a model system consisting of 2*D* membrane patches with defined sizes and reflective boundaries. Single particles were tracked to determine their mean time until they reached the escape opening. We overcame the lifetime limitation of our fluorescent probe by re-scaling the experimentally determined mean first passage time by the fraction of escaped particles *p.* Our approach was confirmed by matching random walk simulations, featuring lifetime-limited and immortal particles. The experimental results of the mean first passage time as a function of the opening coincide with the NEP theory and complementary simulations. We see excellent agreement in both absolute values and systematically concerning the escape opening size *ε*.

## Results and Discussion

### An *in vitro* model system for the systematic testing of the narrow escape problem

For our model system, we use a micro-structured substrate to prepare uniform circular phospholipid membrane patches with a defined diameter. This substrate consisted of circular openings in a continuous gold film deposited on a glass cover slip. These openings were prepared by a colloidal templating approach. In brief, polystyrene microspheres were deposited on a glass substrate, Figure 1 a, and a gold film (100nm) was thermally evaporated, Figure 1 b. The microspheres acted as a shadow mask to prevent gold deposition in areas covered by the particles (39). Upon removal of the microspheres, uniform circular wells surrounded by a continuous gold film remained on the substrate, Figure 1 c-e. We subsequently functionalized the gold with 1-dodecanethiol to passify the surface and prevent the formation of a membrane. Subsequently, small unilamellar vesicles consisting of SOPC (1-stearoyl-2-oleoyl-sn-glycero-3-phosphocholine) were allowed to fuse on the glass substrate within the circular wells, forming well-defined circular supported lipid bilayer (SLB) patches, Figure 1 f. These membrane patches were doped with fluorescently labeled lipids, DOPE-Atto647N (1,2-dioleoyl-sn-glycero-3-phosphoethanolamine with Atto647N label), that diffused laterally in the plane of the SLB.

**Fig. 1.**
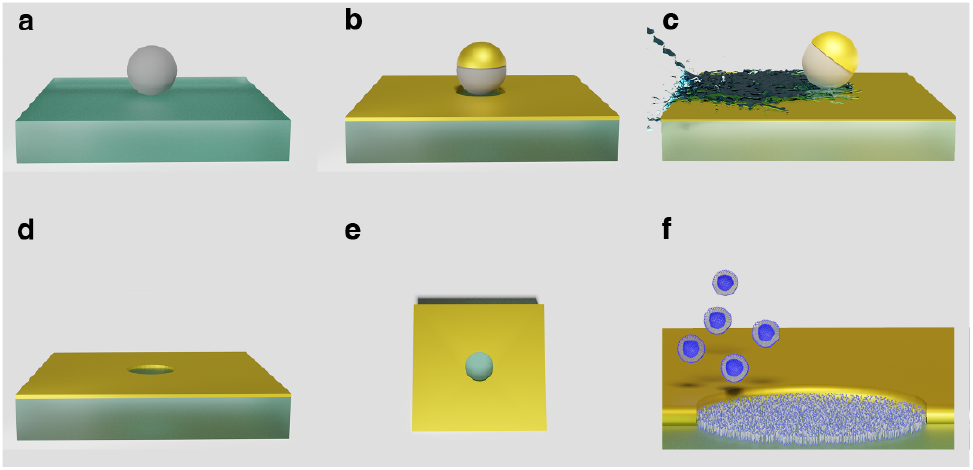
Sketch of the preparation procedural of the mixed colloid-metallization technique. **a** Deposition of the colloids. **b** Metal evaporation of titanium and gold followed by **c** removal of the colloids by rigid washing. **d** Finished scaffolds were cleaned and stored in NTC buffer. **e** Top view and **f** side view of a scaffold/ membrane patch after introduction of a SLB from vesicle fusion.

For the bilayer preparation, imaging and measurements, the substrate was placed in a custom-made observation chamber, which allowed imaging through the transparent glass discs with an inverted microscope. The small diameter of the discs enabled us to probe the full area of the formed membrane patch. Single-molecule localizations were deduced from Gaussian fits, Figure 2 b, on the captured single-emitter intensity profiles, Figure 2 a,c, The radius of the formed membranes was determined by single-molecule fluorescence microscopy (SMFM) via the localization pattern of the fluorescent lipids over time, Figure 2 d. Analyzing 63 domains, we determined the average radius to be 2.44 *μm* with a standard deviation of 0.13 *μm* in agreement with the nominal radius of the beads used to generate the scaffold, *r_bead_* = 2.5*μm*.

**Fig. 2.**
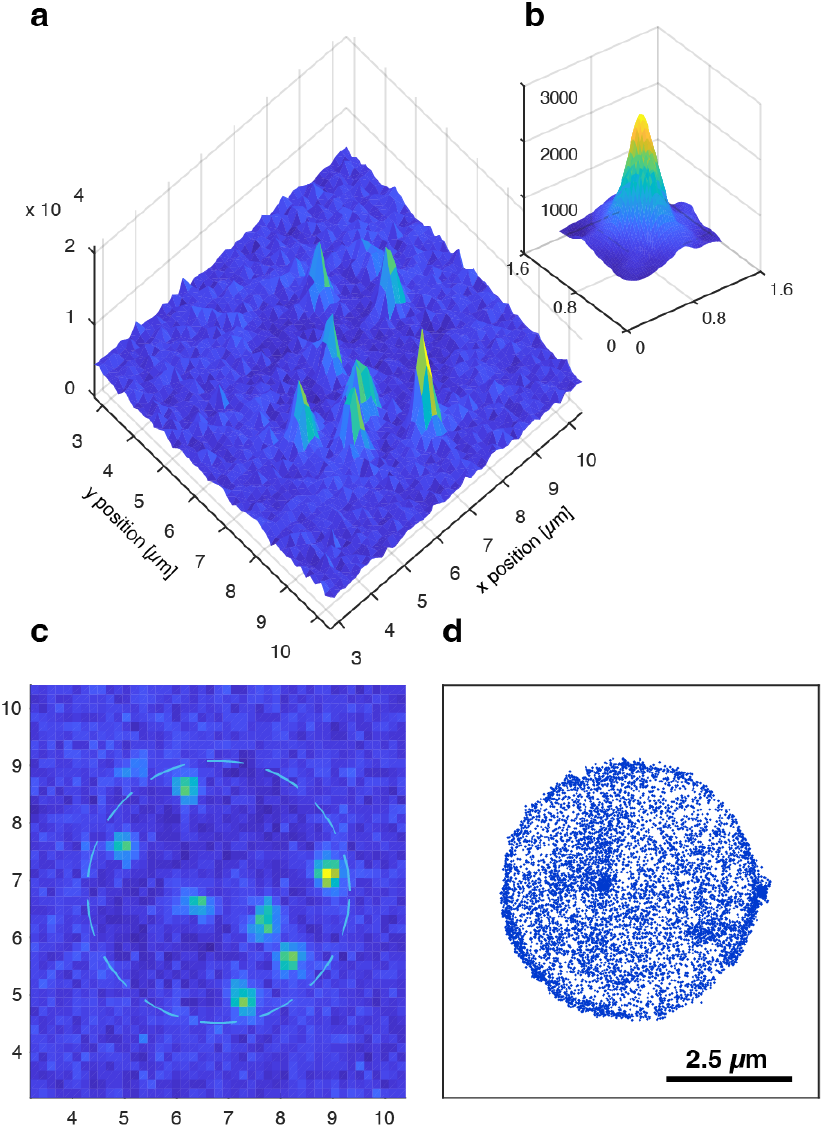
Single-molecule localizations of fluorescent lipids in a circular model membrane. **a** Snapshot of single-molecule intensity profiles. **b** Gaussian fit on the intensity profile of a single emitter. **c** Top view on the single-molecule intensity profiles depicted in a. The dashed circle indicates the rim of the membrane patch. **d** All single-molecule localizations found in an exemplary movie, depicted by dots.

To ensure that our model system was suitable to test the NEP in a disk, we validated the following properties: (i) membrane integrity, (ii) lipid diffusive behavior, and (iii) the reflective properties at the domain boundaries.

#### (i) Membrane integrity

We evaluated the integrity of the membrane patches by calculating the defect area fraction of the membranes. The mean defect area fraction of 63 membrane patches was only 3.6 % with a corresponding median of 2.6 %, Figure SI 3. The linear dependency of the mean squared displacement over time

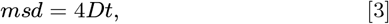

(1)) is a widely accepted measure for free diffusion. Here, only traces with a minimum trace length of 10 steps were analyzed. The *R*^2^ of a linear fit to the msd data yielded 0.98, indicating free diffusion within the membrane patches. The average diffusion coefficient of the tracer lipid was determined to be *D* = 2.29 ± 0.05*μm*^2^/*s* from the slope of the msd over time plot, Figure 3, a. This result matched diffusion coefficients measured in comparable SLB membrane models at room temperature (40, 41).

From the solution of the diffusion equation in one dimension, Equation 5, we know, that normally distributed step lengths are an indication for free diffusion.

#### (ii) Lipid diffusive behavior

To thoroughly check that the lipids in the membrane patches fulfill this criterion for diffusivity, we extracted the distribution of 1D displacements from the measured traces. To exclude steps involved in the reflection at the rim, and thereby eliminate the effects of the confinement, traces were grouped concerning their location at the rim or in the center of the patch. The width of the rim region was chosen to include a large percentage of one-step events while the percentage of two-step events was still reasonably low. The 1D distribution of 13,353 steps (8,349 steps) from the center (rim) region of the domains were found to be normally distributed, 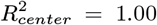, 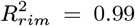, Figure 3 b.) The Gaussian fit of the step length yielded a mean and std of 0.01 ± 0.21*μm* and 0.02 ± 0.15*μm*, for the center and rim, respectively. From these distributions and using Equation 6 we calculated site-specific diffusion coefficients in the center and rim region to complement the average diffusion coefficient determined from the msd analysis. The diffusion coefficient in the center was almost twice as large as at the rim: *D_center_* = 2.21 ± 0.01*μm/s* and *D_rim_* = 1.13 ± 0.02*μm/s*. The reduced diffusion coefficient at the rim points towards an energy loss during reflection. This energy loss at the boundary might be caused by a weak interaction of the lipid probe with the boundaries to the gold film.

**Fig. 3.**
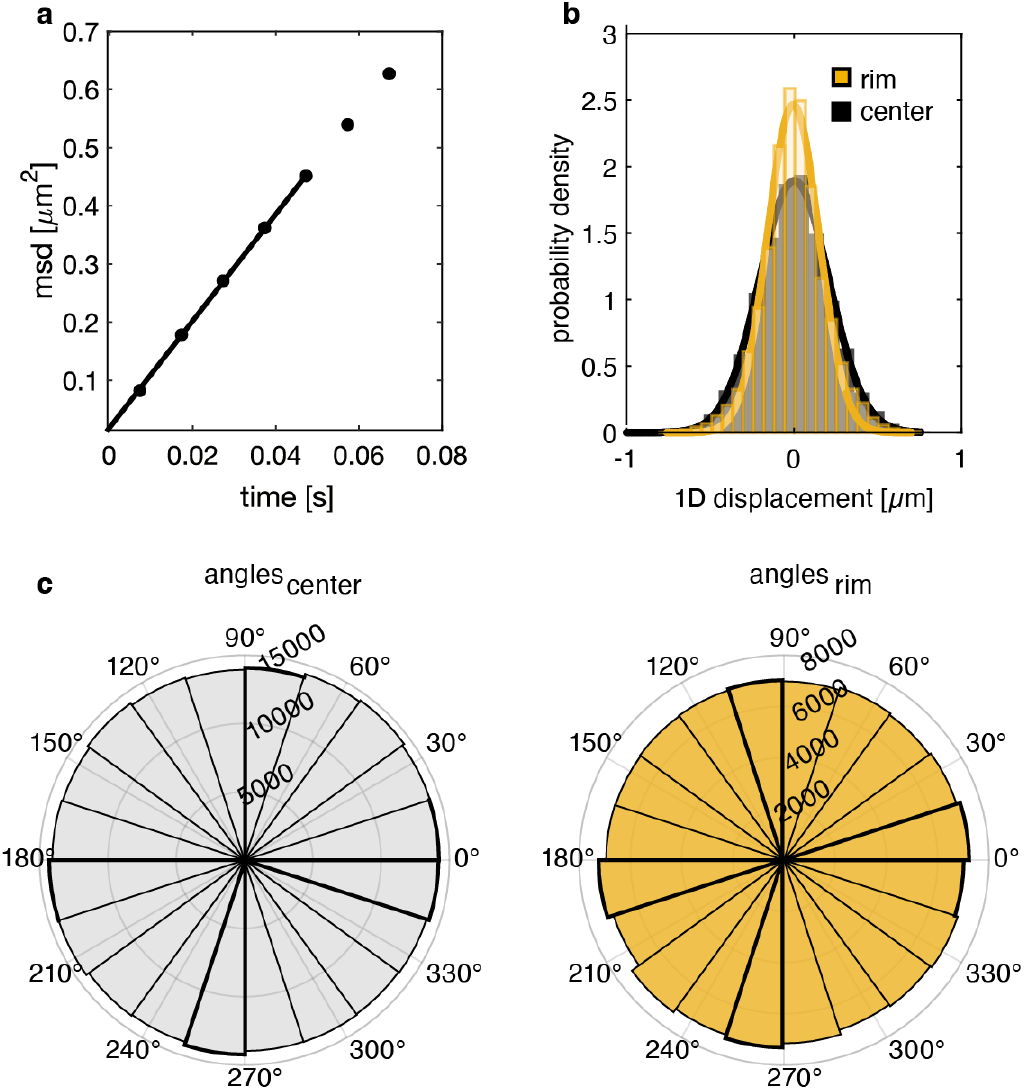
Characterization of the model system. **a** Mean squared displacement over time plot of the tracer lipids in the membrane patch. The error bars display the standard deviation and are of the size of the markers. The line represents a linear fit to the first 5 data points. **b** Distributions of 1D diffusion steps of the tracer lipids in the center (black) and at the rim (yellow) of the domain. The full lines represent Gaussian fits. **c** Distributions of the angles between consecutive steps of the trajectories in the center (left) and rim (right) region. The data shown represents all membrane patches measured.

#### (iii) Reflective properties at the domain boundaries

To evaluate whether the reflection at the boundaries of the membrane patches was biased to the direction, we used the angles between consecutive steps of the lipid trajectories as a measure. The angles were collected from trajectories located at the rim and in the center of the membrane patches separately (details in SI). In both cases the angles were distributed evenly: *α* ∈ [0°, 360°], Figure 3 c). 282,576 and 141,564 angles from the center and rim regions were analyzed, respectively. These reference measurements confirm that our model system fulfilled the requirements for testing the NEP (i - iii).

### Determination of the mean first passage time from lifetime-limited traces

After the positive evaluation of the model system, we experimentally determined the mean first passage times, *τ*, of diffusing lipids for a variety of virtual escape sites at the rim of the diffusion domain and compared it to the theoretical prediction, Equation 2, as well as matching random walk simulations. Single-molecule imaging was performed at 100 *Hz*, and illumination intensity of 1 *kW/cm*^2^. The lipid probes exhibited an ATTO label in the far-red wavelengths spectrum (647N, λ_*abs*_ = 646 nm, λ_*em*_ = 664 nm, extinction coefficient: *ϵ_max_* = 150, 000 *M*^-^1 *cm*^-^1).

The fluorescence signal from individual ATTO647N molecules was localized by 2D Gaussian fitting, 2 b, with a precision of *σ_exp_* = 20 *nm*. This result agreed with the prediction by Thompson et al., Equation 4 (42),

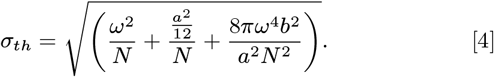

Here, N denotes the number of photons detected per single emitter (N = 379), *ω* the signal width (*ω_exp_* = 336 *nm*), *a* the pixel size (*a_exp_* = 160 *nm*), and *b* the variations in the background (*b_exp_* = 331 *nm*). Using our experimental values given in parenthesis resulted in *σ_th_* = 19 *nm*.

Trajectories were obtained by single-molecule tracking, relying on the principle of maximum probability (43). The total time of each recorded movie was chosen long enough to allow the labeled lipid to probe the full area of the membrane patch. As a result, high coverage with single emitter localizations was achieved. This information allowed us to deduce the boundary of each membrane patch automatically.

To find the escape times of the single emitters, artificial escape openings on the boundary were defined in the analysis. The virtual opening was continuously rotated around the disc, so that the entire boundary was probed over time. With this method, we eliminated bias in choosing the opening, whilst fully exploiting the available microscopy data. The time it took individual lipids to travel to a pre-defined virtual escape window, *ET*, was collected. A fit, Equation 7, to the exponential tail of the probability density histogram of the *ETs*, Figure 4 b, left us with the mean time 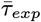 until escape. The mean time 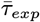 was determined for a systematic variation in escape opening size *ε*, from 1.1 to 12.5 % of the circumference of the membrane patch. The smallest opening was chosen comparable to the size of one pixel in the experiment. The results in Table SI 1 show, that 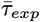 obtained from the collected traces was much shorter than the theoretically predicted mean first passage time. According to Equation 2, we calculated the theoretical mean first passage time, 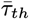, to be ≈ 14 - 8 s (for *ε* ∈ [0.011, 0.125]). However, the lengths of the trajectories collected in the experiments were exponentially distributed with a mean length of only 0.60 s and thus much shorter than 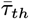. As a consequence, even though ATTO 647N is known for its photostability (34), direct assessment of *τ* was still hampered by the fluorophore lifetime.

**Fig. 4.**
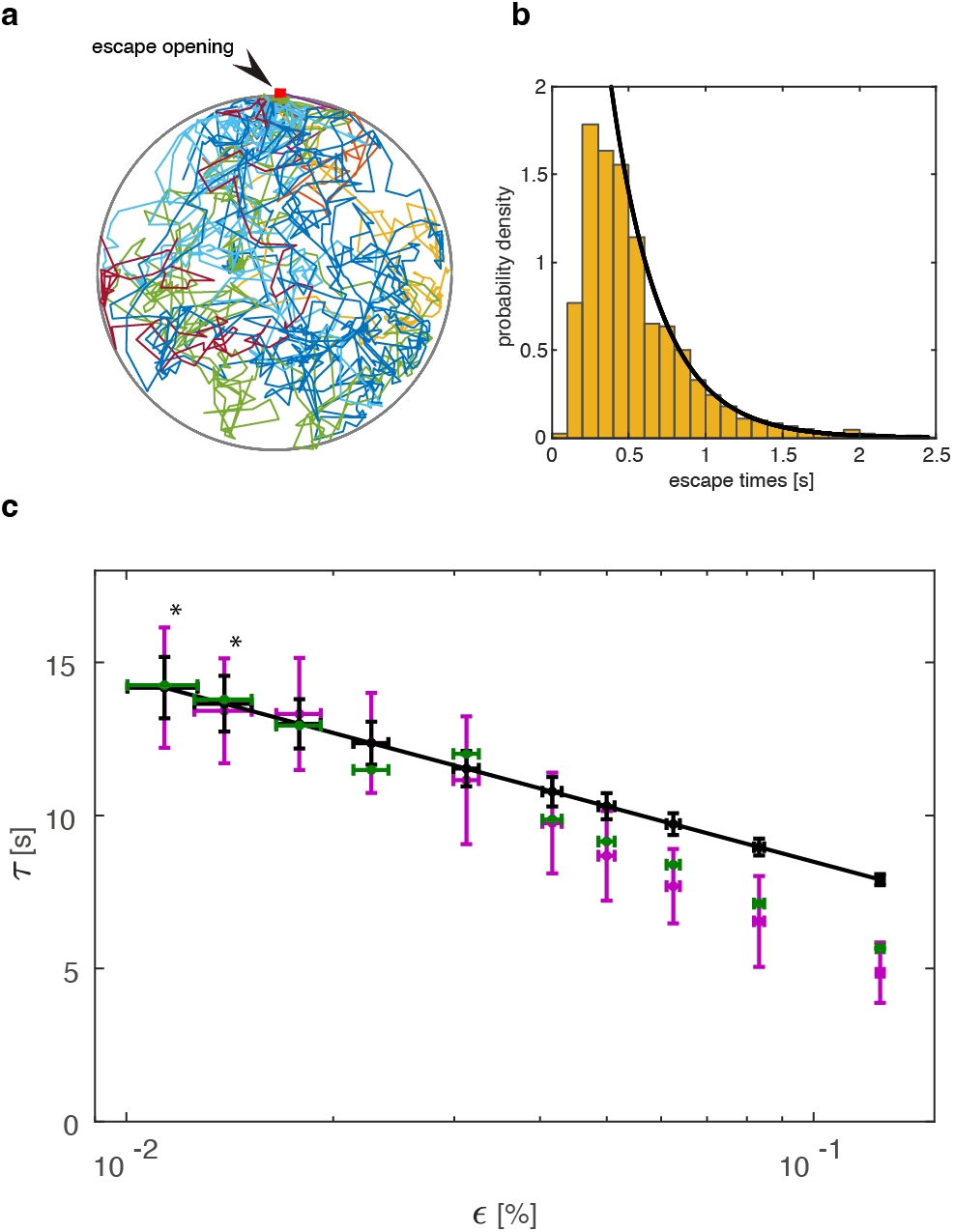
The effect of the escape opening on the mean first passage time determined from experimental traces and simulations. **a** Exemplary single-molecule traces of particles hitting an escape opening (arrow) of *ε* = 1%. **b** Exemplary experimental distribution of escape times for *ε* = 1%. The solid line is the exponential fit to the probability density histogram of the collected escape times. **c** Comparison of the lifetime-corrected experimental mean first passage time with theory and optimized random walk simulations as a function of the escape opening *ε*. The theoretical values, *τ_th_*, are indicated as a black line; black dots highlight *τ_th_* at the escape openings where experiments and simulations were carried out. Lifetime-corrected results of the *in vitro* experiments, *τ_corr, exp_*, are shown in magenta. Corresponding random walk simulations with immortal particles are depicted in green. Simulations, at the most narrow escape openings, were carried out without a localization error and are marked with an asterisk. Vertical error bars of *τ_corr, exp_*, indicate the combined absolute standard error of the fit to all collected *ET*s and *p*s. Vertical error bars of *τ_th_* mark the combined error in *r* and *D.* Vertical error bars of *τ_sim_* represent the error from the fit to all collected escape times. Horizontal error bars are valid for all three data sets and indicate the absolute standard error of *ε*. Additional information on the errors and details on the simulations can be found in the SI and methods sections.

To resolve this discrepancy, caused by lifetime limitation, we considered the actual absorbing probability *p* which can be approximated by the fraction of absorbed particles, here escaped emitters, with respect to all particles present in the domain (see Methods and Equation 8). The absorbing probability *p* depends on the size of *ε*. We found that it varied from 3 to 8 % for all escape openings included. For example, in the case of *ε* = 1.1 % re-scaling of *τ_exp_* = 0.43 s with *p* = 3 % according to Equation 11 resulted in *τ_corr, exp_* = 14.18 s which was in good agreement with *τ_th_* = 14.17 s. All lifetime corrected *τ_exp,corr_* are displayed alongside with *τ_th_* in Table SI 1 and Figure 4.

### Experimentally determined mean first passage times support the theoretical prediction of *τ* ∝ ln(1/*ε*)

After we successfully re-scaled the experimentally determined mean first passage times, we systematically studied the influence of the size of the escape opening site on the mean time until escape. In Figure 4 c our experimental results, *τ_exp,corr_*, are depicted in magenta. They show the expected logarithmic dependence on the window size *ε* ∈ [1.1%, 5%]. However, for escape openings larger than 5 % of the domain circumference, experiment and theory, *τ_th_* depicted in black, gradually diverged: this divergence was expected since *τ_th_* is an approximation valid for *ε* ≪ 1 only. We found that our re-scaled *in vitro* experiments agree well with the theoretical prediction in the limit of small escape openings both in absolute numbers and concerning the logarithmic dependence of the mean first passage time on the escape opening. In the light of this good agreement, we conclude that the observed reduced diffusion coefficient at the rim had a negligible effect on the time *τ_exp,corr_* in the case of a sufficiently large diffusion domain. However, if smaller membrane patches are to be used in future experiments, one should consider introducing additional passivation of the gold to prevent edge effects. For a comparison of experiment and theory over the full range of *ε* see Table SI 1.

### Experiment-inspired simulations of the NEP in 2D confirm the effectivity of the correction factor

To challenge the applicability of the re-scaling factor *p*, we ran random walk simulations with lifetime-limited particles, mimicking the characteristics of the *in vitro* experiments with its uncertainties. To achieve this goal, we created a disc with reflective boundaries, as the simulation environment. The disc featured the mean radius measured in the experiment and an escape site of various sizes. The escape site was enlarged by the localization precision of the single emitters in both dimensions. In the simulation, each Brownian particle diffused with a diffusion coefficient drawn randomly from a normal distribution based on the mean and standard deviation of the measured diffusion coefficient.

The lifetime limitation was introduced by randomly drawing the maximum allowed trace length of a particle from a pool of exponentially distributed values. The pool was chosen to match the average fraction of escaped particles, *p*, in the experiment. Analogs to the analysis of the mean first passage time of experimental traces, the result of the simulation, *τ_sim_*, was re-scaled by *p* to achieve *τ_sim, corr_*.

As a reference, simulations were run with immortal particles. These particles were allocated a trace length that allowed them to always reach the escape site *ε* (See Methods for a detailed description of the simulations). The lifetime-corrected mean first passage time *τ_sim,corr_* matched the results from traces not restricted in lifetime *τ*_*sim*, ∞_ see Figure SI 4 cyan and green data points, respectively. Thus, the concept of re-scaling, motivated by theoretical considerations was successfully applied to experimental data and was additionally demonstrated *in silico*, see Figure SI 4 for a full comparison of the different simulation scenarios.

### Experiment-inspired simulations of the NEP in 2D show agreement with the *in vitro* experiment

For a comprehensive anal-ysis of the narrow escape problem in a disc, we compared our experimental result for the mean first passage time with the random walk simulations featuring immortal particles introduced above. Figure 4 c summarizes the results of the *in silico* and *in vitro* experiments, in green and magenta, respectively. The experimentally-inspired random walk simulations presented here agree with the NEP theory for small escape openings (*ε* < 4%) as well as with experimental results in the full range of escape openings studied.

In summary, we introduced a high-quality scalable model system for systematic testing of the narrow escape theory in a disc. As stated above, the mean first passage time can be unambiguously determined by three parameters: the domain area, *A*, the diffusion coefficient, *D*, and the relative escape opening *ε*. We used a comprehensive approach integrating theory, simulation, and experiment to prove the logarithmic dependence of 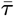 on the relative escape opening.

Our model system offers the excellent opportunity to additionally address the functional dependency of 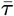 on the remaining two parameters *A* and *D*. The domain area can be easily varied with the size of the colloids used in the preparation of the membrane patches while alternate temperature settings will directly modulate the diffusion coefficient of the tracer lipids. However, the range for *A* is limited by the energy loss during the reflection event (lower bound) and the traceability of the chosen label (upper bound). The lower bound may be further pushed by an additional passivization step to the protocol. Possible temperature settings to control the diffusion coefficient are constrained by the phase behavior of the lipid matrix and the achievable acquisition frequency. The use of optimized labels like SeTau-647-NHS (44) (quantum yield Φ = 0.61, extinction coefficient *ϵ_max_* = 200, 000 *M*^-^1 *cm*^-^1)(45) or Janelia Fluor-JF646 (quantum yield Φ = 0.54, extinction coefficient *ϵ_max_* = 152, 000 *M*^-^1 *cm*^-^1) (46), which provide enhanced brightness and photo-stability will extend the testable range of both parameters (*A* and *D*).

If we now consider a specific biological application in immunology or parasitology, the diffusion domains will take unique, complex geometries. Ergo, it becomes difficult to find a suitable theoretical model to calculate 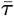 analytically. In these cases, 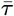 must be determined experimentally. However, the sample properties and technical limitations generally do not allow the full observation of rare escape events. Here, our approach to dealing with lifetime-limited traces provides a straightforward solution to directly measure 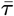 in those scenarios that were experimentally inaccessible so far.

## Materials and Methods

### Preparation of glass cover slips

Hydrophilic properties and clean-liness of the microscopy coverslips were ensured using the following protocol. Glass coverslips 22×22 of a controlled thickness of 0.17 ± 0.01*mm* (Karl Hecht, Sondheim (GER)) were plasma cleaned with ambient air (Expanded Plasma Cleaner 230 V, Harrick Plasma (NY)) at a high power setting (30 W) for 20 minutes. The glass was rinsed with double deionized water (ddH2O) and consecutively sonicated (10 min at 37 kHz), in acetone, ethanol, and methanol, respectively. Every time the solvent for the sonification was changed, the glass coverslips were washed extensively with ddH2O. Subsequently coverslips were immersed in RCA (Radio Corporation of America) solution (1:1:5, ammonia: hydrogen peroxide: ddH2O) for 45 min at 80°C. The cleaning protocol was completed by the last washing step with at least 1 L ddH2O. Clean and hydrophilic glasses were stored in ddH2O and used within three days.

### Microstructuring of glass cover slips

A colloid lithography approach was used to create the defined microstructuration of the glass substrate. To this end, polystyrene microspheres (*r_bead_*=2·5 *μm*; Thermofisher) were used to template the glass discs in a continuous gold film. A droplet of dispersion of the microspheres (0.1 wt-percent) was added to the precleaned glass coverslip and rapidly dried using a nitrogen stream to randomly distribute the microspheres on the substrate. Subsequently, a thin layer of titanium (2nm, used as an adhesion promoter) was thermally evaporated (Torr International Inc, THE3-KW) at normal incidence onto the microsphere-covered coverslip, directly followed by the thermal evaporation of a gold layer of 100 *nm.* The polystyrene microspheres were removed by rinsing with ddH2O to produce circular glass discs within a continuous gold film.

### Lipid work

Model membranes (micro-structured SLBs) were formed by vesicle fusion. Small unilamellar vesicles (SUVs) were prepared from SOPC (1-stearoyl-2-oleoyl-sn-glycero-3-phosphocholine) or a mixture of SOPC and DOPE ATTO 647N (1,2-dioleoyl-sn-glycero-3-phosphoethanolamine). Both lipids: Avanti Polar Lipids, Alabaster (AL). Lipids dissolved in chloroform were dried under a nitrogen stream, rehydrated in NTC buffer at a final lipid concentration of 1 mg/ml, and vortexed briefly. The following sonication (37 kHz) of the NTC-lipid-mix was conducted for 10 min in pulsed mode followed by 10-30 min in sweep mode. Whilst sonication, the temperature was kept below 20 °*C* using ice in water bath. Membrane patches were formed by incubation of SOPC SUVs on micro-structured glass for at least 1h at 37 °*C*. Trace amounts of fluorescently labeled lipids were introduced into the SLB via incubation with SUVs prepared from SOPC and 0.005 mol % DOPE ATTO 647N labeled lipids. Usually, only a few *μ*l of a 1 mg/ml SUV solution were sufficient. The molecular models of the used lipids are displayed in Figure SI 5. Excess vesicles were removed by washing with 20 ml of NTC. Imaging was done at 25 °*C*. For the preparation of the SLB and imaging, the scaffold was placed in a custom-made observation chamber. The latter allowed imaging through the glass bottom of the scaffold with an inverted microscope. The average size of a lipid headgroup is 0.7 *nm*^2^ and the discs have a mean area of 18.7 ^2^ (47). Therefore, we estimate a total of 27×10^6^ SOPC molecules per disc, including around 10 labeled DOPE molecules. Thus approximately 0.3 ppm of lipids are labeled.

Lipid stocks were stored dry at −80°*C* for long terms. For short-term storage, lipids were solved in chloroform (HPLC grade, Carl Roth, Karlsruhe (GER)) and stored at −20 °*C*.

### Single-molecule imaging

Single-molecule imaging data was obtained with a home-build microscopy setup. Here, an inverted wide-field microscope (Leica DMI 6000B, Leica) was equipped with a 515 *nm* laser line (Cobolt, Fandango, 50 mW) and a 640 *nm* line (Cobolt, MLD, 100 mW), an AOTF (AOTF nc-400.650-TN, AA opto-electronic, Orsay), and a high numerical aperture objective (HCX PL APO 100x/ 1.47 OIL CORR TIRF, Zeiss). Signals were detected by the back-illuminated EMCCD-camera (iXon3 DU-897, Andor Technology, Belfast) using appropriate filter combinations (zt405/ 514/ 633rpc (dichroic, Chroma, Rockingham) and 550/49 BrightLine HC (emission filter, Semrock, Rochester)). Movies of up to 10000 frames were taken in crop mode at 100 Hz with a frame size of 95 x 95 px (image pixel size: 160 nm) and laser intensity of 1 kW/cm^2^). Synchronized sample illumination and camera readout were ensured by using the camera-ready signal as a trigger for AOTF opening at a suitable frequency. Image acquisition was controlled by the open-source software *μ*Manager 1.4 (48).

### Single-molecule localization and tracking

Localization and tracking of single fluorescently labeled lipids were conducted with MATLAB routines as previously described in (43, 49). Image data from preparations that did not yield an integer and thus fluent membrane, enabling free diffusion, were excluded from the analysis. In brief, single-molecule intensity distributions were fitted with a 2D Gaussian function to determine their localization in x and y as well as their width and integrated intensity. The resulting list of positions was further filtered with predefined characteristics of Atto647N under the given illumination conditions. The remaining positions served as the input for the tracking algorithm. A probabilistic algorithm was used to connect the single-molecule positions in adjacent frames of a movie. Assuming free diffusion, a translational matrix was built up to calculate the probabilities of all possible connections. Subsequently, trajectories were connected by maximizing the total probability of all connections. In solid-supported lipid bilayers on micro-patterned glass, a precision of ~ 20 *nm* was achieved for ATTO 647N labeled lipids. For details see (49).

### Determination ofthe diffusion coefficient from the distribution of 1D step lengths

The solution of the diffusion equation for one dimension yields the probability *ρ* for finding a particle at distance *l* after time *t* (49):

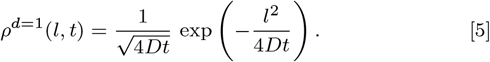

We determined the 1D step length distributions in the center and at the rim of the domain and fitted them with normal distributions.1D steps are obtained from decomposition of the 2D displacement of the particles in the x- and y-components. For the definition of the rim and center region see SI. From the fitted standard deviation, *std*, the diffusion coefficient in the center and at the rim was calculated according to:

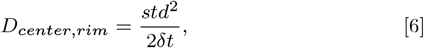

where *δt* was the time lag in between two consecutive frames of a movie.

### Reflectivity test

The reflectivity of the domain boundaries was assessed from the distribution of angles between consecutive diffusion steps in the center of the domain versus the rim. The width of the rim region to be included in the analysis was chosen to comprise all possible step lengths up to 80 % of the cumulative step length distribution calculated according to Equation 5 using the previously determined average diffusion coefficient of 2.29 *μm*^2^/s. The angles were computed by basic trigonometry.

### Analysis of single-molecule trajectories to determine the individual escape times and the mean first passage time

To test the narrow escape theory, Equation2, only traces that started at least from a distance of *r* were used in the mean first passage time analysis. To meet this condition, first, the actual size of the membrane domains was determined from trajectories. Further, the outline and center of mass of all individual membrane patches were found. Second, the outline was partitioned into sections of desired size. Each section was designated to be the escape site once. Last, we sorted for trajectories that did start at least *r* from the escape site. The time until the escape of every single lipid (*ET*) was calculated from the number of steps performed by the particle until colocalization with the escape site. Both the parts of the traces before and after hitting the escape site were used for the analysis if the endpoint was also at least *r* away from the escape site. The distribution of trace lengths of the resulting trace parts was in a range of 10-261 steps, with a mean trace length 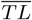 of 22 steps. The individual escape times from all particles, escape sites, and movies were collected and plotted in a probability density histogram. The mean first passage time was determined from the fit in the exponential tail of the histogram:

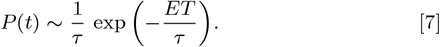

The tail region was found automatically by maximizing the corresponding *R*^2^ value. An example is shown in Figure 4 b.

### Estimating the escape time from incomplete trajectories

The experimental approach to measure the mean first passage time *τ* was limited by the lifetime and traceability of the used fluorescent emitter, here ATTO 647N. During the time of observation *T*, the trajectories disappeared randomly. Consequently, the total number of trajectories *n_T_* recorded per domain was much larger than the number of absorbed trajectories *n_a_*. The absorbing probability *p* which represents the probability that a trajectory is absorbed before it disappears can be computed from the fraction of absorbed trajectories. The absorbing probability is defined and approximated by the formula:

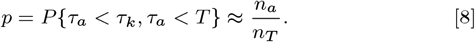

We used this absorbing probability to derive the relation between the theoretical narrow escape time 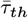 and the 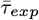 estimated from experimental trajectories. The absorbing probability *p* provides a correction factor for the time 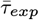, which is relevant especially if the mean duration of the trajectories is much shorter than the absorbing time 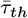, as observed here.

To derive the relations between these parameters, we recall that the mean first passage time 〈*τ_d_*〉 for a particle that disappears has two components: one due to absorption at the narrow window and the second is due to killing or bleaching. Thus, since the two events are independent, using Bayes’law, we obtain:

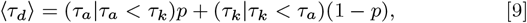

where *τ_a_* and *τ_k_* is the first absorbing and killing time, respectively. We assume that the end time *T* is sufficiently long and neglect the remaining particles that could not be absorbed or killed after time *T*. This result is independent of the killing law for trajectories (20). When the killing time is much smaller than the absorbing time, 〈*τ_k_*|*τ_k_* < *τ_a_*)〉 ≪ 〈*τ_a_*|*τ_a_* < *τ_k_*〉, we neglect the second term, in Equation 9 and thus we are left with the relation 〈*τ_th_*〉 = 〈*τ_a_*|*τ_a_* < *τ^k^*〉 where the total disappearance time 〈*τ_d_*〉 can be measured experimentally:

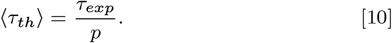

We determined the correction factor *p* empirically: both for experimental *p_exp_*, and simulated *p_sim_* cases using relation Equation 8. In each case, the absorbing probability *p* depends on the size of the escape window *ε* (20). Thus, we calculated *p*(*ε*) for all analyzed escape window sizes. In the experimental case, we collected the fraction of escaped particles with respect to the number of all particles for each *ε* from all 63 movies and 1/*ε* escape windows, separately. From the resulting Weibull distribution, we determined the mean 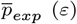 by least-square fitting. This 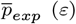 was then used to re-scale *τ_exp_* according to the identity:

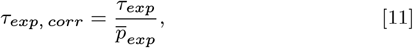

where *τ_exp, corr_* being the corrected and *τ_exp_* the measured mean first passage time, respectively. In case of simulations with lifetime-limited traces, we found *p_sim_* simply from the ratio of the number of escaped particles and the total number of simulated traces.

### In silico experiments

*In silico* experiments were carried out with parameters (*D_exp_* = 2.29 ± 0.05*μm*^2^/*s*, *r_exp_* = 2.44 ± 0.13*μm*) matching the *in vitro* experiments. For the simulation of trace_*i*_, *D_i_* and *r_i_* were drawn independently from normal distributions. Dimensionless particles were simulated to perform a random walk in a confined domain until escape through a predefined window of size *a* ± *σ* on top of the domain. The spatial uncertainty *σ* was derived from the *in vitro* experiments. When interacting with the boundaries of the domain, the particles were relocated to their last position before encountering the boundary. For each step, a random angle was chosen *α* ∈ [0, 2*π*] while the step lengths *l* fulfilled the criterion for the probability *ρ*, Equation 12 (49):

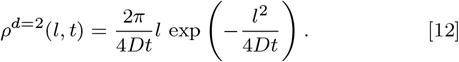

where *D* was the diffusion coefficient and *t* the time lag. Particles were counted as escaped when hitting a predefined window *a*. The time until escape, *ET*, was collected. The mean first passage time of simulated traces, *τ_sim_*, was determined from a fit, Equation 7, to the exponential tail of the distribution of the collected *ET*s as described above for the experimental data.

At large relative escape windows *ε* the time step of the simulation was chosen to be *δt* = 10 *ms*, as in the experiment. With decreasing *ε*, the average diffusion step 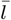 became of the order of the escape opening *a*, resulting in a systematic overestimation of *τ*. As a consequence *δt* was adjusted to fulfill the criterion of 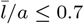, Table SI 1. In simulations at very small escape openings, where the value of the spatial uncertainty *σ* amounted to at least 10 % of the opening *a* we did see that *τ_sim_* seemed to level off. Similar findings were made for simulations with lifetime limitations, which were executed to test *p*. As a consequence, we conducted simulations with ideal localization precision (*σ* = 0) for escape openings, where 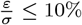 applied. This was the case for *ε* ∈ [0.014, 0.011]. At this optimized spatial resolution, flattening of the *τ* (*ε*) curve was prevented and the experimental observation, as well as the theoretical result, was recovered.

Simulations with and without lifetime limitations were carried out to test the performance of the correction factor. The latter was realized in simulations with extremely long trace lengths to ensure the escape of all particles (*τ*_*sim*, ∞_). To mimic the experimental situation, simulations with lifetime limitations were carried out (*τ_sim, p_*). This was achieved by simulation of traces with exponentially distributed lengths and a mean equal to the mean step length found *in vitro*. The resulting *τ_sim, p_* was corrected as described for the microscopy data *τ_sim, corr_* = *τ_sim, p_/p_sim_*.

### Statistics and Error calculation

*τ_th_* was calculated according to Equation 10. In order to ensure comparability with the experimental result, *D_exp_* and *r_exp_* were used as found in the experiment - including the associated uncertainties *SE_D_* and *SE_r_*. *SE_D_* and *SE_r_* were then classically propagated to achieve *SE_τ_th__*. For the *in vitro* experiment, the total error of *τ_corr_* was calculated from the sum of relative errors of the two contributing parameters (*τ_exp_* and *p* see Equation 11) :

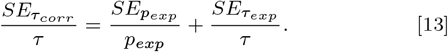

Since the experimental data (*τ_exp_*) is a result of a fit to the distribution of individual *ET*s, *SE_τ,exp_* represents the standard error of the fit parameter. *p_exp_* and its corresponding standard error, 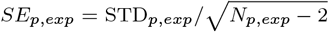 were obtained from the mean (*p_exp_*) and standard deviation (STD_*p,exp*_) of a Weibull distribution of all (N_*p,exp*_) collected *p_exp_*.

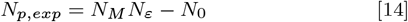

depending on the number of movies *N_M_*, the number of escape windows in each movie *N_ε_* (*N_ε_* = 1/*ε*), and the number of escape sites with no registered escapes *N*ø.

For the *in silico* experiments, the uncertainties in *D_exp_* and *r_exp_* were already included in the simulations. Thus, the standard error in the fit on the exponential tail of the *ET* distribution is represented in the vertical error bar.

The error in *ε* = *a*/ (2*r_exp_π*) is displayed as a horizontal error bar in the *τ* vs *ε* plots. This error has two contributions, the error in the radius of the domain and the localization precision of the single emitter:

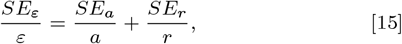

with *SE_a_* being the localization precision *σ* of the single particles.

## Data availability

The data sets generated during and/or analyzed during the current study are available from the corresponding author upon reasonable request.

## Code availability

Information on custom code used in this project is available from the corresponding author upon reasonable request.

## Materials & Correspondence

Correspondence and material requests should be addressed to E.M.

## ACKNOWLEDGMENTS

This work was funded by the Deutsche Forschungsgemeinschaft, project number: 391332795 (S.F.F) and grant number VO 1824/6-2 (N.V.). We thank Markus Engstler for fruitful discussions. We thank the precision mechanics workshop of the Biocenter for expert technical assistance.

## References

1. Einstein A (1905) Über die von der molekularkinetischen theorie der wärme geforderte bewegung von in ruhenden flüssigkeiten suspendierten teilchen. Annalen der physik 4.

2. Sokolov IM (2019) Extreme fluctuation dominance in biology: On the usefulness of wastefulness: Comment on” redundancy principle and the role of extreme statistics in molecular and cellular biology” by z. schuss, k. basnayake and d. holcman. Physics of life reviews 28:88–91.

3. Singer A, Schuss Z, Holcman D (2006) Narrow escape, part ii: The circular disk. Journal of statistical physics 122(3):465–489.

4. Lagache T, Holcman D (2017) Extended narrow escape with many windows for analyzing viral entry into the cell nucleus. Journal of Statistical Physics 166(2):244–266.

5. Lagache T, Holcman D (2008) Quantifying intermittent transport in cell cytoplasm. Physical Review E 77(3):030901.

6. Lagache T, Dauty E, Holcman D (2009) Physical principles and models describing intracellular virus particle dynamics. Current opinion in microbiology 12(4):439–445.

7. Reingruber J, Holcman D (2011) The narrow escape problem in a flat cylindrical microdomain with application to diffusion in the synaptic cleft. Multiscale Modeling & Simulation 9(2):793–816.

8. Engstler M, et al. (2004) Kinetics of endocytosis and recycling of the gpi-anchored variant surface glycoprotein in trypanosoma brucei. Journal of cell science 117(7):1105–1115.

9. Bruce D,, et al. (1897) Further report on the tsetse fly disease or nagana, in zululand. Harrison & Sons.

10. Cross GA (1975) Identification, purification and properties of clone-specific glycoprotein antigens constituting the surface coat of trypanosoma brucei. Parasitology 71(3):393–417.

11. Vickerman K (1969) On the surface coat and flagellar adhesion in trypanosomes. Journal of cell science 5(1):163–193.

12. Horn D (2014) Antigenic variation in african trypanosomes. Molecular and biochemical parasitology 195(2):123–129.

13. Hartel AJW, et al. (2015) The molecular size of the extra-membrane domain influences the diffusion of the gpi-anchored vsg on the trypanosome plasma membrane. Scientific reports 5(1):1–12.

14. Florimond C, et al. (2015) Bilbo1 is a scaffold protein of the flagellar pocket collar in the pathogen trypanosoma brucei. PLoS Pathog 11(3):e1004654.

15. Holcman D, Schuss Z (2004) Escape through a small opening: receptor trafficking in a synaptic membrane. Journal of Statistical Physics 117(5):975–1014.

16. Schuss Z, Singer A, Holcman D (2007) The narrow escape problem for diffusion in cellular microdomains. Proceedings of the National Academy of Sciences 104(41):16098–16103.

17. Pillay S, Ward MJ, Peirce A, Kolokolnikov T (2010) An asymptotic analysis of the mean first passage time for narrow escape problems: Part i: Two-dimensional domains. Multiscale Modeling & Simulation 8(3):803–835.

18. Cheviakov AF, Ward MJ, Straube R (2010) An asymptotic analysis of the mean first passage time for narrow escape problems: Part ii: The sphere. Multiscale Modeling & Simulation 8(3):836–870.

19. Grebenkov DS, Holcman D, Metzler R (2020) Preface: new trends in first-passage methods and applications in the life sciences and engineering. Journal of Physics A: Mathematical and Theoretical 53(19):190301.

20. Holcman D, Schuss Z (2015) Stochastic narrow escape in molecular and cellular biology. Analysis and Applications. Springer, New York 48:108–112.

21. Holcman D, Schuss Z (2013) Control of flux by narrow passages and hidden targets in cellular biology. Reports on Progress in Physics 76(7):074601.

22. Pedone D, Langecker M, Abstreiter G, Rant U (2011) A pore-cavity-pore device to trap and investigate single nanoparticles and dna molecules in a femtoliter compartment: confined diffusion and narrow escape. Nano letters 11(4):1561–1567.

23. Pomp W (2017) Ph.D. thesis (Leiden University).

24. Mathai PP, Liddle JA, Stavis SM (2016) Optical tracking of nanoscale particles in microscale environments. Applied Physics Reviews 3(1):011105.

25. Lévy R, Shaheen U, Cesbron Y, Sée V (2010) Gold nanoparticles delivery in mammalian live cells: a critical review. Nano Reviews 1(1):4889. PMID: 22110850.

26. Suarez-Kelly LP, et al. (2021) Antibody conjugation of fluorescent nanodiamonds for targeted innate immune cell activation. ACS applied nano materials 4(3):3122–3139.

27. Garifo S, Stanicki D, Ayata G, Muller RN, Laurent S (2021) Nanodiamonds as nanomaterial for biomedical field. Frontiers of Materials Science pp. 1–18.

28. Murase K, et al. (2004) Ultrafine membrane compartments for molecular diffusion as revealed by single molecule techniques. Biophysical Journal 86(6):4075–4093.

29. Gemeinhardt A, et al. (2018) Label-free imaging of single proteins secreted from living cells via iscat microscopy. JoVE (Journal of Visualized Experiments). (141):e58486.

30. Shimomura O, Johnson FH, Saiga Y (1962) Extraction, purification and properties of aequorin, a bioluminescent protein from the luminous hydromedusan, aequorea. Journal of Cellular and Comparative Physiology 59(3):223–239.

31. Yuste R (2005) Fluorescence microscopy today. Nature methods 2(12):902–904.

32. Snapp EL (2009) Fluorescent proteins: a cell biologist’s user guide. Trends in Cell Biology 19(11):649–655. Special Issue – Imaging Cell Biology.

33. Croce AC, Bottiroli G (2014) Autofluorescence spectroscopy and imaging: a tool for biomedical research and diagnosis. European journal of histochemistry: EJH 58(4).

34. Clausen MP, Lagerholm CB (2011) The probe rules in single particle tracking. Current Protein and Peptide Science 12(8):699–713.

35. Holcman D, Marchewka A, Schuss Z (2005) Survival probability of diffusion with trapping in cellular neurobiology. Physical Review E 72(3):031910.

36. Holcman D (2007) Modeling dna and virus trafficking in the cell cytoplasm. Journal of Statistical Physics 127(3):471–494.

37. Meerson B, Redner S (2015) Mortality, redundancy, and diversity in stochastic search. Physical review letters 114(19):198101.

38. Grebenkov DS, Rupprecht JF (2017) The escape problem for mortal walkers. The Journal of chemical physics 146(8):084106.

39. Bley K, et al. (2018) Hierarchical design of metal micro/nanohole array films optimizes transparency and haze factor. Advanced Functional Materials 28(13):1706965.

40. Rose M, Hirmiz N, Moran-Mirabal JM, Fradin C (2015) Lipid diffusion in supported lipid bilayers: a comparison between line-scanning fluorescence correlation spectroscopy and single-particle tracking. Membranes 5(4):702–721.

41. Fenz SF, Merkel R, Sengupta K (2009) Diffusion and intermembrane distance: case study of avidin and e-cadherin mediated adhesion. Langmuir 25(2):1074–1085.

42. Thompson RE, Larson DR, Webb WW (2002) Precise nanometer localization analysis for individual fluorescent probes. Biophysical journal 82(5):2775–2783.

43. Schmidt T, Schütz G, Baumgartner W, Gruber H, Schindler H (1996) Imaging of single molecule diffusion. Proceedings of the National Academy of Sciences 93(7):2926–2929.

44. Patsenker L, et al. (2008) Fluorescent probes and labels for biomedical applications. Annals of the New York Academy of Sciences 1130(1):179.

45. Kudriavtseva YO, et al. (year?) Setau dyes next generation long-wavelength biomedical labels with advanced characteristics.

46. Grimm JB, Brown TA, English BP, Lionnet T, Lavis LD (2017) Synthesis of janelia fluor halotag and snap-tag ligands and their use in cellular imaging experiments in Super-Resolution Microscopy. (Springer), pp. 179–188.

47. Smaby JM, Muderhwa JM, Brockman HL (1994) Is lateral phase separation required for fatty acid to stimulate lipases in a phosphatidylcholine interface? Biochemistry 33(7):1915–1922.

48. Edelstein A, Amodaj N, Hoover K, Vale R, Stuurman N (2010) Computer control of microscopes using *μ*manager. Current protocols in molecular biology 92(1):14–20.

49. Fenz SF, Pezzarossa A, Schmidt T (2012) The Basics and Potential of Single-Molecule Tracking in Cellular Biophysics, Comprehensive Biophysics. (Oxford: Academic Press) Vol. 2.

