## Supplemental Information for "Experiments in micro-patterned model membranes support the narrow escape theory"

### Supplemental Data

**Determination of rim and center regions.** To analyze the traces in the rim and the center area of the membrane patches the corresponding areas needed to be defined. We decided on using the probability density function of possible step lengths as a measure. Random 1-step and 2-step events were generated based on experimental diffusion coefficient and time lag. We compared the two distributions, see Figure SI 1, to find a threshold in units px to which extent the rim was taken. We found, that for a rim of 2 px ($320nm$) the majority (71 %) of possible step lengths had a higher probability to belong to the 1-step events than to the 2-step events. Further, this was still a threshold within which as few 2-step events as possible were included in the rim area (46 %).

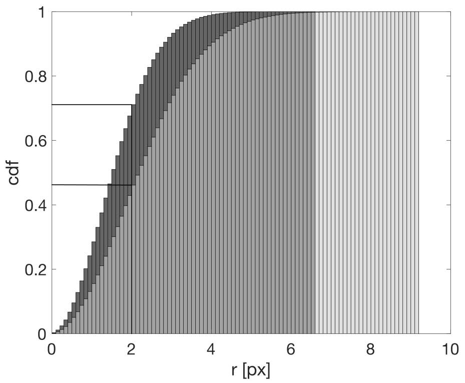


**Fig. SI 1. Comparison of the probability density functions of random 1-step and 2-step events based on experimental data.** The cumulative probability of a step from a 1-step event (dark) or a 2-step event (light) to travel a distance of r in the membrane domain. The black line is marking the thresh hold, set at a 2 pixel travel range.

**Test for Weibull distribution of the correction factors $p$ from the individual escape windows.** For every movie, a test of Weibull distribution was performed on the distribution of $p$ for each window of a certain size separately. The test was performed with an inbuilt function of MATLAB (wblplot.m). It plots the data against a theoretical Weibull distribution in such a way that a real Weibull distribution will collapse to a line. It was found that the $p$'s corresponding to the individual escape openings were indeed Weibull distributed. Here, examples of the test on the data Figure SI 2a and the Weibull fit on the $p$ data Figure SI 2b are shown. The exemplary figures SI 2a, b correspond to data from all movies and $\varepsilon = 1\%$.

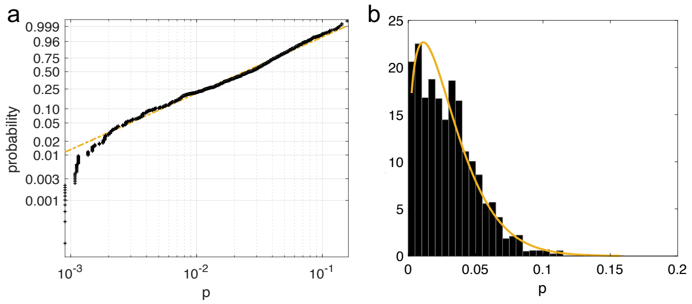


**Fig. SI 2. Confirmation of Weibull distribution and fit to the collected $p$. a** depicts the Weibull cumulative probabilities of the determined correction factors. Agreement of the data (black) with the guideline (yellow) is indicating a Weibull distribution. **b** shows an exemplary distribution of $p$ (black) fitted with a Weibull fit (yellow).

**Membrane integrity test.** Only an intact membrane facilitates diffusion. Immobile particles indicate a membrane defect. That is why we calculated the area fraction covered by immobile particles as a measure for membrane quality. It was calculated from the ratio of pixels occupied by immobile and mobile particles, respectively. Both behaviors were mutually exclusive - no pixel fell in both categories. Immobile particles were classified according to the squared displacement enclosing the full trace. We sorted the squared displacement of all traces in ascending order and observed a plateau followed by a steep increase. The turning point, 4.08 $px^2$, was chosen as a threshold to separate immobile and mobile traces. We observed a clear separation of traces exhibiting free long-range diffusion and those performing confined diffusion. Figure SI 3 shows, the distribution of the mean defect areas of all individual membrane domains. The mean and median defect fraction was found to be only 3.6 % and 2.6 %, respectively.

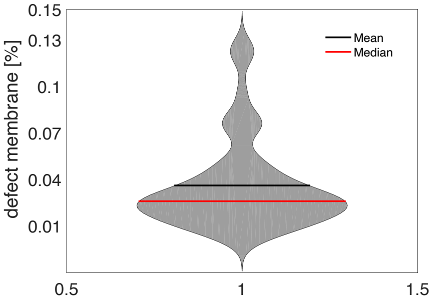


**Fig. SI 3. Violin Plot.** Violin plot of the fraction of defect membrane surface for all 63 membrane patches.

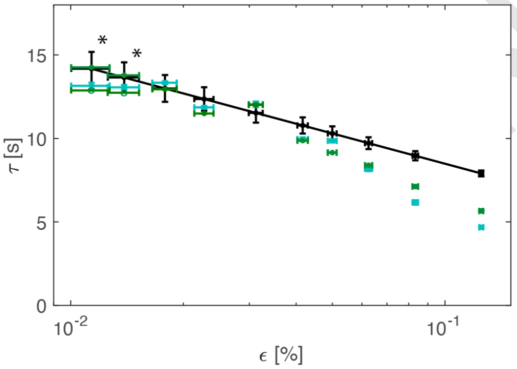


**Fig. SI 4. Effectivity of the correction factor in simulations.** Comparison of the mean first passage time, $\tau$, determined from theory (black line) and simulations for a range of escape openings, $\varepsilon$. Experiment-inspired random walk simulations with and without lifetime limitation are depicted in cyan and a variation of green, respectively. Filled markers in green marked with an asterisk are from simulations using a localization precision of $\sigma = 0$. Corresponding empty markers indicate simulations with $\sigma = 20nm$, which were not used in the final comparison, but illustrate the leveling effect. For a description of the errors see Figure 4.

**Table SI 1. Overview of simulation parameters and all explicit values for $\tau$ from experiments, simulations and theory. From left to right: relative escape opening, temporal resolution in the simulation $\delta t[ms]$, ratio $\bar{l}/a[\%]$ of the mean step length $\bar{l}$ in simulations, and the window size a, $\bar{\tau}_{sim,\infty}$ from simulations with infinite particle lifetime, $\bar{\tau}_{sim,corr}$ from simulations with lifetime limited particles, the uncorrected measured mean first passage time $\bar{\tau}_{exp}$, the theoretical mean first passage time $\tau_{th}$, the lifetime corrected $\bar{\tau}_{exp,corr}$ and $p_{exp}$, which is the correction factor for the experimental data. Values marked with an asterisk are from simulations using a localization precision of $\sigma = 0$. All $\bar{\tau}$ are in $[s]$.**

| $\varepsilon$ | $\delta t[ms]$ | $\bar{l}/a[\%]$ | $\bar{\tau}_{sim,\infty}$ | $\bar{\tau}_{sim,corr}$ | $\bar{\tau}_{exp}$ | $\tau_{th}$ | $\bar{\tau}_{exp,corr}$ | $p_{exp}$ |
|---|---|---|---|---|---|---|---|---|
| 1/8 | 10 | 16 | 5.66 | 4.68 | 0.411 | 7.90 | 4.86 | 0.084 |
| 1/12 | 10 | 23 | 7.12 | 6.17 | 0.432 | 8.96 | 6.54 | 0.066 |
| 1/16 | 10 | 31 | 8.40 | 8.19 | 0.435 | 9.72 | 7.69 | 0.057 |
| 1/20 | 10 | 39 | 9.15 | 9.85 | 0.438 | 10.30 | 8.68 | 0.050 |
| 1/24 | 10 | 47 | 9.87 | 9.96 | 0.455 | 10.78 | 9.75 | 0.047 |
| 1/32 | 10 | 63 | 12.00 | 12.05 | 0.464 | 11.53 | 11.15 | 0.042 |
| 1/44 | 5 | 60 | 11.49 | 11.86 | 0.464 | 12.36 | 12.37 | 0.038 |
| 1/56 | 4 | 70 | 12.94 | 13.33 | 0.466 | 12.99 | 13.31 | 0.035 |
| 1/72 | 2 | 65 | $13.78*$,12.73 | 13.06 | 0.435 | 13.65 | 13.42 | 0.032 |
| 1/88 | 1.5 | 69 | $14.26*$,12.88 | 13.15 | 0.434 | 14.17 | 14.18 | 0.031 |

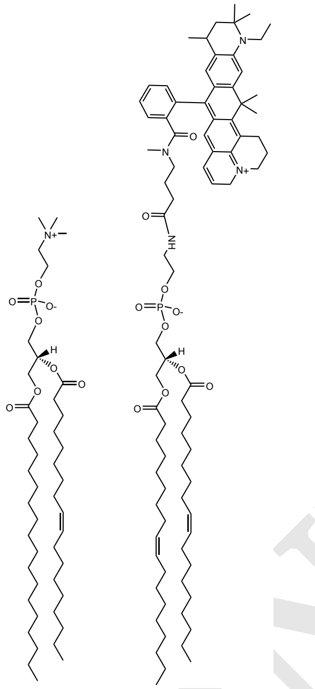

SOPC DOPE-ATTO647N

**Fig. SI 5. Structures of the used lipids.** The molecular models of SOPC (1-stearoyl-2-oleoyl-sn-glycero-3-phosphocholine) and DOPE ATTO 647N (1,2-dioleoyl-sn-glycero-3-phosphoethanolamine).

PREPRINT
